# Plant litter chemistry and associated changes in microbial decomposition under drought

**DOI:** 10.1101/2025.11.12.688006

**Authors:** Brian Chung, Shi Wang, Zhao Hao, Steven D. Allison, Ashish A. Malik

## Abstract

Drought has consequences for microbial decomposition rates, including indirect effects through changes in plant litter chemistry. Here we studied the impact of a decade-long drought on plant litter chemistry and microbial decomposition traits in a semi-arid ecosystem during an 18-month litter bag experiment. We investigated litter sourced from four conditions: grass and shrub vegetation under ambient and reduced precipitation. We hypothesized that litter chemistry drives microbial decomposition capabilities and enzyme activity, either due to vegetation differences or drought effects on litter chemistry. Fourier Transform Infrared Spectroscopy was used to characterize litter chemistry; we found that carbohydrate-rich grass litter decomposed faster than more recalcitrant shrub litter which was richer in lignin and lipids. There were significant changes in litter chemistry under drought but no increase in lignin fraction suggesting that drought does not make litter more recalcitrant. Metagenomics-derived decomposition genes and extracellular enzyme activity were higher in grass litter; patterns related to differences in substrate supply. Genes linked to lignin depolymerization decreased in abundance under drought. However, most decomposition genes and enzyme activities were not significantly affected by drought thereby maintaining decomposition rates. Microbial community succession with higher abundance of fungi at early and bacteria at later stages of decomposition corresponded with genes for fungal and bacterial necromass recycling along with protein accumulation over time. We demonstrate minimal litter chemistry-mediated effects of drought but show significant changes in community composition and their decomposition capabilities over time highlighting that complex microbial-chemical interactions under climate change can influence ecosystem-scale processes.

**Importance:** Climate change is causing more severe and frequent droughts in semi-arid ecosystems, affecting soil microbes breaking down plant litter. Our research focusses on understanding the less studied pathway of drought impact on microbes via changes in plant litter chemistry. Drought can alter the plant litter chemistry, by changing the composition and physiology of plants, which can alter microbial decomposition and ecosystem-level carbon cycling. We investigated litter decomposition traits of microbial communities in grass and shrub litter under long-term drought. There were significant changes in litter chemistry under drought but no increase in lignin fraction. Despite this, microbial communities maintained their decomposition capabilities under drought highlighting ability of microbes to adapt and continue functioning. We also demonstrate unique microbial community succession patterns and dead biomass recycling which can have implications for carbon cycling rates in the ecosystem. This study sheds light on the complex microbial interactions that affect ecosystem functioning under climate change.

## Introduction

Changes in precipitation regimes are projected to occur with climate change, with increasing drought having already been observed in semi-arid ecosystems across the globe such as in California, western South America, and the Mediterranean (1). Increasing evapotranspiration due to an increase in the co-occurrence of high temperatures and low precipitation (2) is leading to more severe droughts. Furthermore, the number of extreme precipitation events, including periods of low precipitation, are also projected to increase (3). In addition, changes in precipitation are not uniform at a regional scale (1), with arid and semi-arid ecosystems in southern California experiencing decreasing annual precipitation (4).

The increasing severity and frequency of droughts can have major effects on key ecosystem processes such as decomposition and the organisms driving it, mainly fungi and bacteria. The effects on decomposer microorganisms can be both direct – through changes in abundance of individual taxa, community composition, and traits for drought stress tolerance – and indirect – through changes in the plant litter chemistry (5–7). Microbial response to drought may not always be consistent and may vary across ecosystems (8) or climate histories (9, 10). Drought in semi-arid ecosystems decreases litter decomposition rates in some systems (11–13) but not others (14). Decreases in decomposition have been attributed to decreases in microbial biomass (13) and the efficiency of extracellular enzymes (15). While drought shifted the overall microbial community composition of an oak forest towards fungal dominance, drought increased both bacterial and fungal abundance in a mixed pine-oak forest such that fungal:bacterial ratios remained unchanged (14). Drought can also shift investment in microbial traits such that allocation toward stress tolerance traits reduces decomposition capabilities.

Litter chemistry can be a major control on decomposition (16, 17), including decomposition in grasslands (18). Drought has been shown to alter litter chemistry (13, 19, 20) through changes in plant physiology (20) and changes in plant community composition (21, 22). These changes in litter chemistry, in turn, affect the microbial community, altering its composition by decreasing bacterial abundance (13), and decreasing investment in extracellular enzyme activity by decreasing proportions of certain litter substrates (15). Therefore, drought can exert direct and indirect effects on litter decomposition, and the indirect effects remain understudied (7).

Here we investigate the effects of a decade-long drought on the decomposition traits of microbial communities during an 18-month litter bag experiment in a semi-arid ecosystem, specifically focusing on microbial resource acquisition traits as influenced by the vegetation type and the corresponding litter chemistry. We tested the impact of drought on plant litter derived from grass and shrub vegetation that experienced either ambient or reduced precipitation for 10 years. We hypothesized that shrub litter decays more slowly than grass litter because it contains more lignin/lipids and less cellulose/hemicellulose while drought alters the chemistry of both litter types, making them decay more slowly. Further, we hypothesized that litter chemistry drives microbial decomposition capabilities and enzyme activity, whether due to vegetation differences or drought effects on chemistry. We tested these hypotheses using Fourier Transform Infrared Spectroscopy (FTIR) to study changes in litter chemistry, shotgun metagenomics to measure decomposition capabilities, and fluorometric extracellular enzyme assays to quantity decomposer activity. We synthesize the knowledge to present evidence of the less-studied effects of drought via changes in plant litter chemistry on microbe-mediated decomposition rates, a key ecosystem function.

## Results

### Shrub litter is more recalcitrant than grass litter

Litter chemistry differed between the vegetation types (Figure 1). The carbohydrate ester spectral areas (Figure 1h, 1i, S1) are likely associated with hemicellulose and pectins that contain ester groups (23). While shrub litter had higher spectral area of carbohydrate C-O stretching (Figure 1e), grassland litter had higher spectral areas of other spectral ranges associated with carbohydrates (Figure 1d, 1h, 1i), indicating that grassland litter had higher overall carbohydrate content than shrub litter. Shrub litter had higher spectral area associated with C-H methyl and methylene deformation (Figure 1b) and lipid C=O stretching (Figure 1c). The spectral range 1450-1475 cm^-1^ has been associated with lignin (24). The spectral range 1700-1750 cm^-1^ has been associated with C=O stretching in ketones and carboxylic acids (23), indicating that this range might be associated with lipids. Shrub compared to grass litter likely had higher proportions of more recalcitrant compounds, namely lignin and lipids (Figure 1b, 1c), indicating that shrub litter is more recalcitrant than grassland litter, consistent with our hypothesized difference in decay rates between these two litter types. Differences in lignin and carbohydrates between the vegetation types are also consistent with other studies that compared one or two litter species from each of the same vegetation types (25, 26). In our study, chemical recalcitrance likely plays the most important role in determining rates of decomposition.

**Figure 1.**
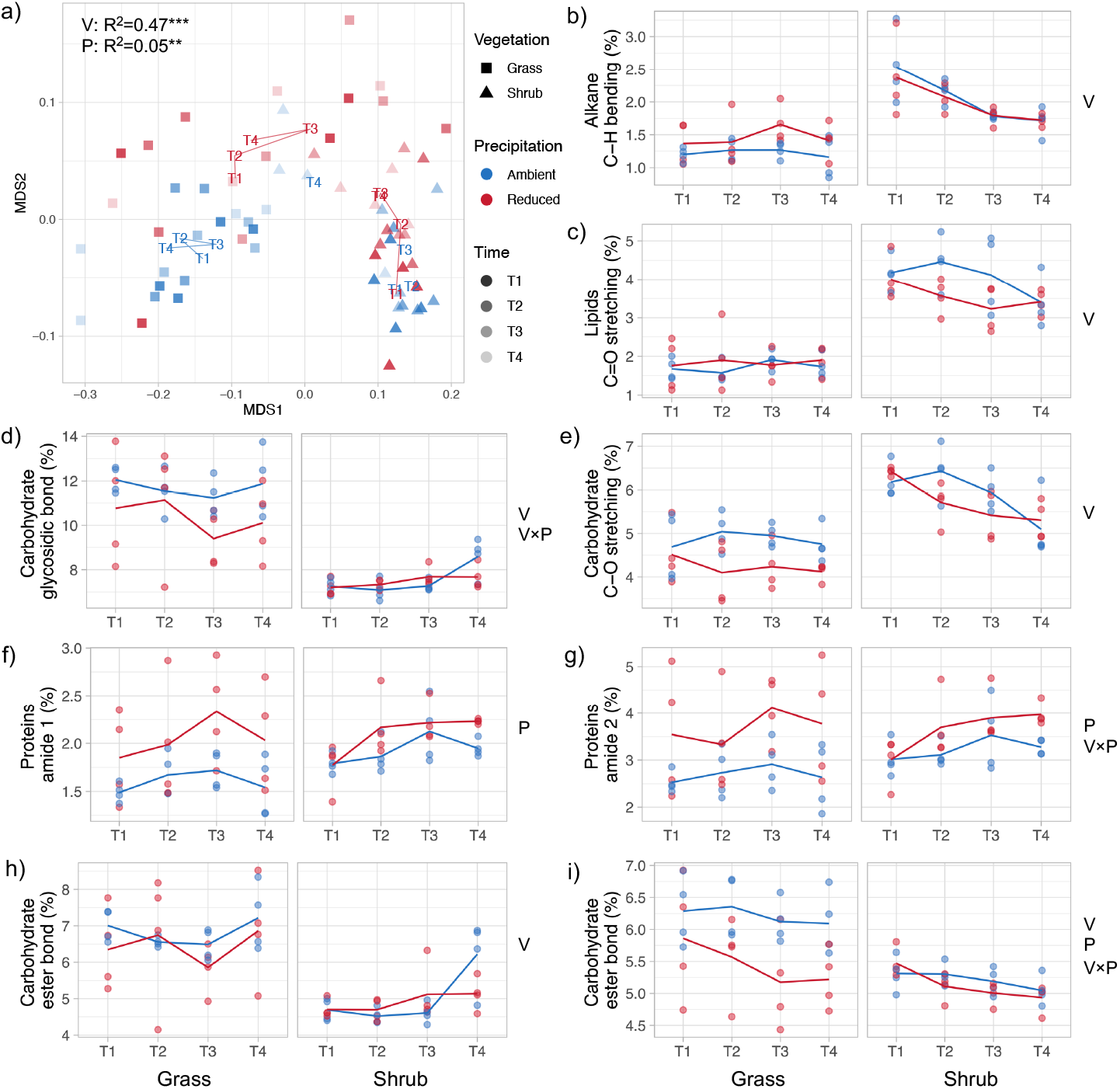
Litter chemistry differences across vegetation and precipitation treatments: a) an NMDS plot of litter chemical composition derived from FTIR with PERMANOVA R^2^ and asterisks indicating significant vegetation (V) or precipitation (P) treatment effect (^***^ *P*<0.001, ^**^ *P*<0.01, ^*^ *P*<0.05). b-i) changes in key compound classes over time for the four litter types; y-axis representing the proportional abundance estimated as the area under the curve assigned to a specific FTIR spectral range with the letters at the side only showing significant vegetation (V), precipitation (P) or interaction effect derived using linear mixed effects models and Tukey’s post-hoc test. b) 1450 - 1475 cm^-1^, C-H deformations in methyl and methylene groups; c) 1700 - 1750 cm^-1^, lipids; d) 1015 - 1080 cm^-1^, C-O deformation of glycosidic bonds; e) 1160 - 1230 cm^- 1^, carbohydrate C-O stretches; f) 1620 - 1645 cm^-1^, amide 1; g) 1545 - 1600 cm^-1^, amide 2; h) 970 - 1015 cm^-1^, carbohydrate ester; i) 1100 - 1160 cm^-1^, C-O stretches in carbohydrate esters.

### Significant changes in litter chemistry with drought

While drought significantly affected litter chemistry of both litter types as shown by an overall precipitation effect on some spectral bands (Table S1), drought had much stronger effects on grass litter than shrub litter (Figure 1). Significant interactions between vegetation type and precipitation were present for carbohydrate glycosidic bonds and a carbohydrate ester spectral range (Figure 1d, 1i, Table S1), with drought only lowering spectral areas in these two ranges in grassland litter (Figure 1d, 1i). The effects of drought on carbohydrates in grassland litter were consistent with decreases in cellulose and hemicellulose that have previously been found in the grassland drought plots of this field experiment (13). Drought did not affect lignin or lipids in either litter type (Figure 1b-c, Table S1), indicating that drought did not increase the recalcitrance of either litter types. This is inconsistent with the effects of drought predicted by our hypothesis.

Amide spectral ranges, which are indicative of proteins, increased over time (Figure 1f-g), consistent with increases of protein in litter over time in other systems (27). Drought increased protein concentration in litter of both types, although the effects were stronger for grass litter than shrub litter (Figure 1f-g, Table S1). Increases in nitrogen under drought have previously been observed elsewhere (19, 20), including in a previous study in the grassland vegetation of this field experiment (13).

### Decomposition genes not strongly affected by drought

The abundance of carbohydrate active enzymes or CAZyme genes for metabolizing carbohydrates – hemicellulose and oligosaccharides – was higher in grass litter than shrub litter (Figure 2d, 2h) while CAZyme genes for lignin were more abundant in shrub litter (Figure 2g). These differences in CAZyme gene abundance, when comparing broadly between the two litter types, are consistent with the differences in litter chemistry. The results across precipitation treatments do not support the indirect effects of drought through plant litter chemistry changes that we hypothesized. CAZyme genes for carbohydrates – cellulose, hemicellulose, starch, polysaccharides, and oligosaccharides – were not affected by drought in either ecosystem (Figure 2c-f, 2h, Table S1) whether drought decreased carbohydrate fractions – as in grassland litter – or not – as in shrub litter (Figure 1d, 1i, Table S1). Lignin-related genes decreased in abundance under drought across both systems (Figure 2g) despite drought not affecting lignin fractions in litter (Figure 1b, Table S1). While these results do not necessarily preclude indirect effects of drought on lignin genes that we hypothesized, they could indicate a direct effect of drought on lignin genes.

**Figure 2.**
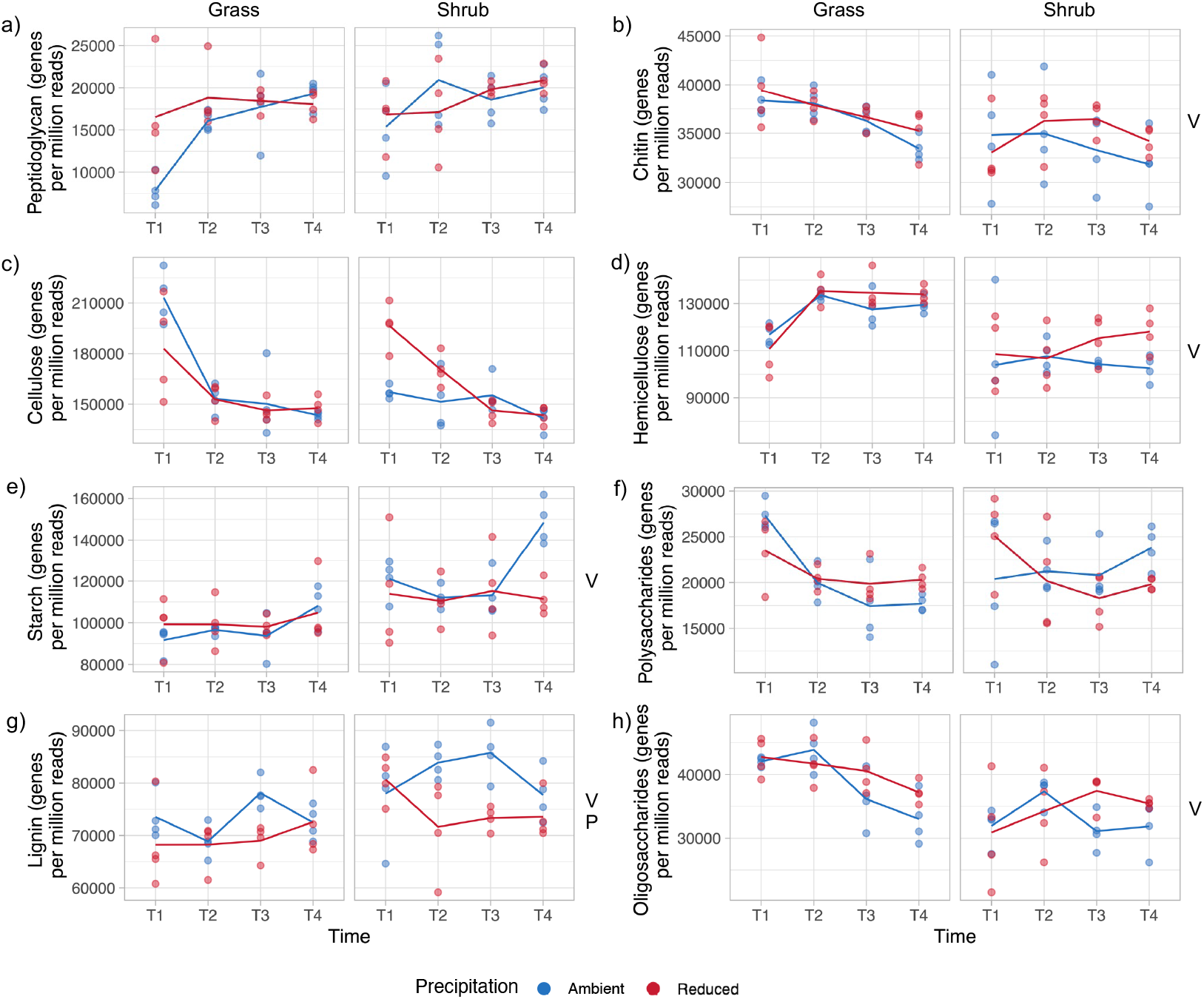
Gene-level decomposition capabilities across vegetation and precipitation treatments: CAZyme gene abundance for putative substrates. a) peptidoglycan, b) chitin, c) cellulose, d) hemicellulose, e) starch, f) polysaccharides, g) lignin, h) oligosaccharides. The letters at the side only showing significant vegetation (V), precipitation (P) or interaction effect derived using linear mixed effects models and Tukey’s post-hoc test.

### Patterns of community succession with decomposition

CAZyme gene abundances related to cellulose, polysaccharides, and oligosaccharides decreased over time (Figure 2c, 2f, 2h) while CAZyme gene abundances related to hemicellulose and starch increased over time (Figure 2d-e), indicating a succession of the decomposition of different substrates. These chemical changes were also linked to changes in microbial community composition over time across both litter types with grass litter experiencing stronger changes (Figure 3). Taxonomic diversity increased over time in both systems (Figure 3a) while fungal:bacterial ratios decreased over time in both systems (Figure 3b). These changes in composition also corresponded with temporal trends in CAZyme genes involved in microbial cell wall metabolism. Gene abundance for peptidoglycan metabolism increased over time (Figure 2a) while chitin genes decreased over time in both systems (Figure 2b).

**Figure 3.**
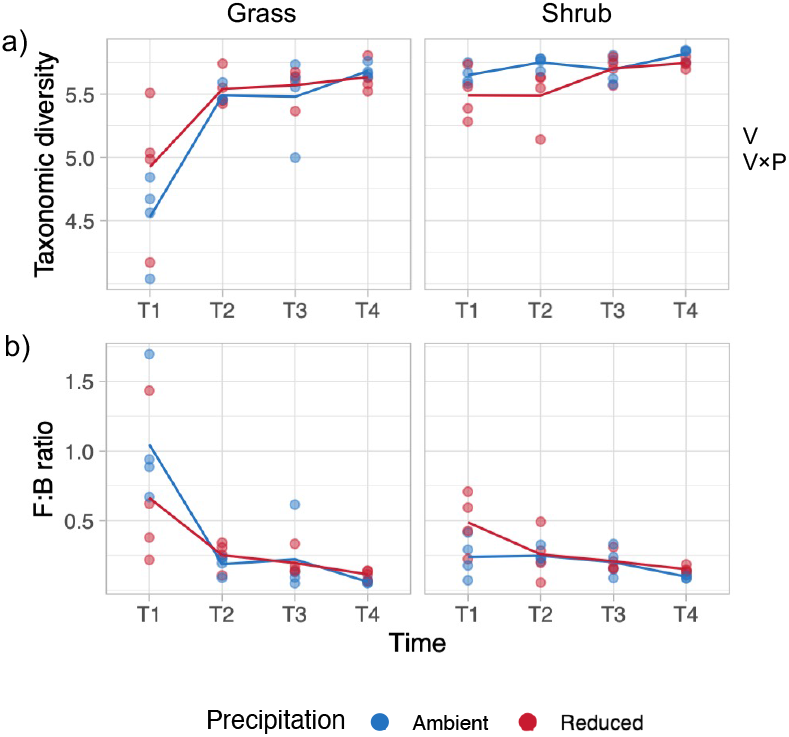
Diversity across vegetation and precipitation treatments: a) Taxonomic diversity presented as alpha diversity based on genus-level annotations derived from metagenomics reads. (b) Fungal:bacterial ratios estimated as read abundance ratios of the two groups from the same dataset. The letters at the side only showing significant vegetation (V), precipitation (P) or interaction effect derived using linear mixed effects models and Tukey’s post-hoc test.

### Extracellular enzyme activity is driven by substrate supply

Enzyme activity tended to be higher in grassland than shrub litter (Figure 4), with statistically significant differences for the enzymes cellobiohydrolase and N-acetyl-β-D-glucosaminidase (Figure 4d, 4f, Table S1) and insignificant differences for ɑ-glucosidase, β-glucosidase, and β-xylosidase (Figure 4a-c, Table S1). The higher carbohydrate content of grassland litter (Figure 1d, 1h, 1i) corresponded with larger pools of extracellular enzymes that target carbohydrates (Figure 4a-d) suggesting that enzyme activity is driven by substrate supply rather than microbial demand.

**Figure 4.**
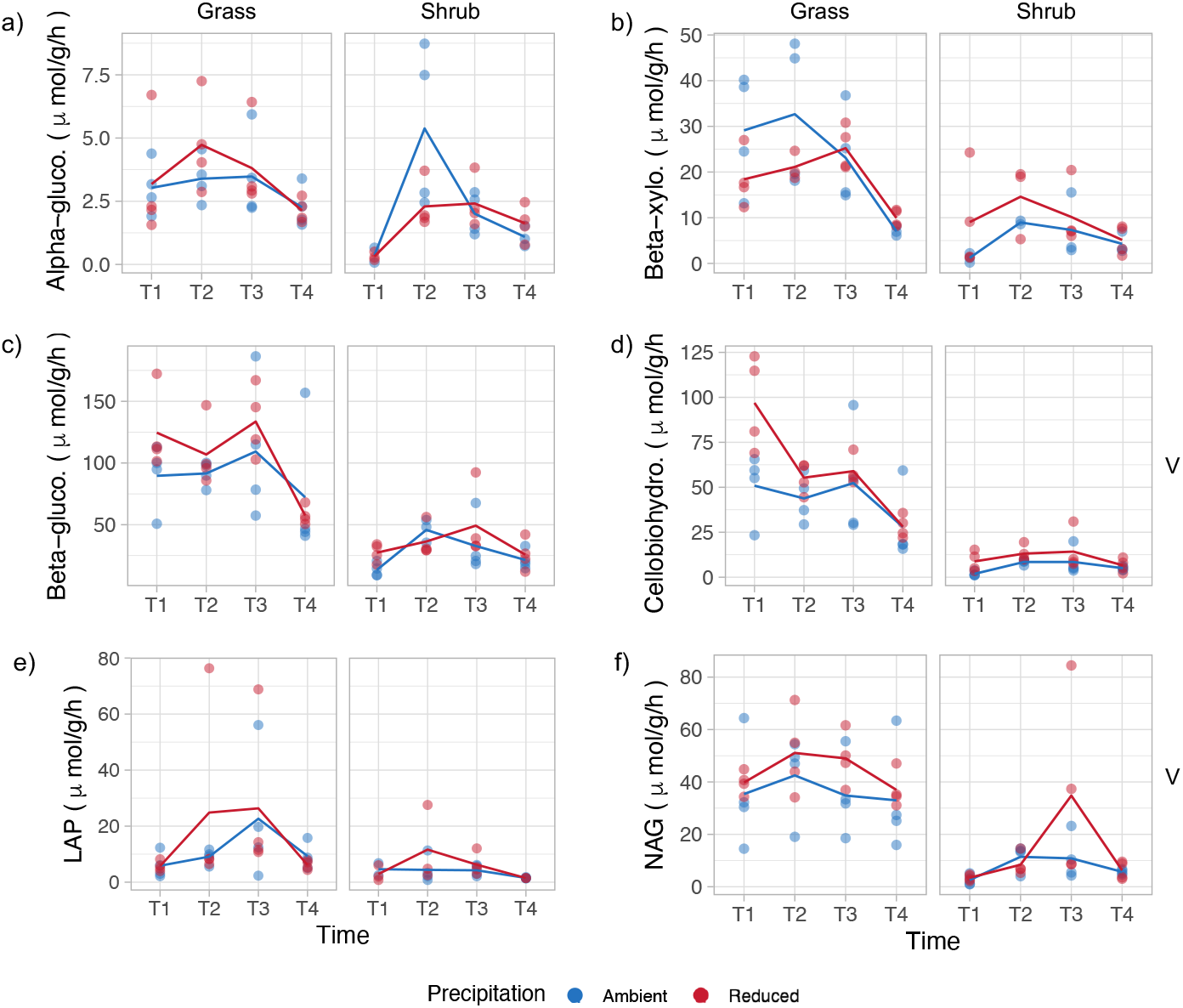
Extracellular enzyme activity measured as enzyme V_max_ for a) ɑ-glucosidase, b) β-xylosidase, c) β-glucosidase, d) cellobiohydrolase, e) leucine aminopeptidase, and f) N-acetyl-β-glucosaminidase. The letters at the side only showing significant vegetation (V), precipitation (P) or interaction effect derived using linear mixed effects models and Tukey’s post-hoc test.

Drought had no statistically significant effect on the activity of any enzymes, either as a main effect or as an interaction with vegetation (Figure 4, Table S1). There was also very high variability across replicates which could have obscured treatment effects. This result does not support the indirect effect of drought on enzyme activity that we hypothesized, as activity of carbohydrate enzymes remained unchanged under drought (Figure 4a-d) whether carbohydrate fractions decreased – as in grassland litter – or remain unchanged – as in shrub litter (Figure 1d, 1h, 1i).

## Discussion

### Induction of enzymes by their substrates

Resource acquisition traits and litter chemistry differed between vegetation types as predicted by our hypothesis. We observed higher CAZyme gene abundance and carbohydrate-degrading enzyme activity in grass litter than in shrub litter (Figures 2d, 2h, 4a-d), which likely explained the faster decomposition rates of grassland litter observed at this site (28). Because grassland litter tended to have higher carbohydrate content than shrub litter (Figure 1d, 1h, 1i), these results are consistent with the theory of induction of enzymes by their substrates (29, 30) and positive associations between CAZyme genes and their substrates that have been observed elsewhere (31, 32). Higher carbohydrate-degrading enzyme V_max_ in the grassland community (Figure 4a-d) could stem from differences in microbial genomic content, with the grassland microbial community’s higher abundance of hemicellulose and oligosaccharide CAZyme genes than the shrub community (Figure 2d, 2h).

Grass litter had lower proportions of recalcitrant compounds such as lignin than shrub litter (Figure 1b), which could also allow for greater enzyme V_max_ in grass litter (33). Microbes that specialize in lignin degradation possess more genes that function in cell signaling pathways rather than hydrolytic enzymes (34). Lignin also adsorbs hydrolases (35, 36), likely decreasing enzymatic breakdown and reducing concentrations of substrates and intermediate degradation products that induce enzyme production (29). Both factors could be further contributing to the gaps in V_max_ between the shrub and grass litter microbial communities.

### Microbial succession and necromass recycling

Along with broad differences in functional gene abundances between the litter communities, we also observed changes that correspond with succession in microbial communities as decomposition progressed. Fungal-bacterial ratios decreased with time (Figure 3b), corresponding with increasing peptidoglycan gene abundance (Figure 2a) and decreasing chitin gene abundance (Figure 2b), indicating microbial communities depolymerizing fungal and bacterial necromass as a carbon source. Some of the peptidoglycan genes we observed are used by bacteria to recycle their cell walls in the process of cell growth (37) which could also explain the increase over time. Decreasing chitin gene abundance is consistent with decreasing abundance of bacteria that decompose fungal cell walls (38) as well as decreasing fungal abundance. Decreasing fungal abundance also corresponded with decreasing trends over time of β-glucosidase, β-xylosidase, and cellobiohydrolase Vmax (Figure 4b-d), trends that have been observed in a temperate oak forest (38, 39). Some studies show that fungi are the main producers of extracellular enzymes (40, 41), and a previous study in our grassland system found that the most abundant fungal taxa explained more variation in extracellular enzyme activity than the most abundant bacterial taxa (42).

### Effects of drought through litter chemistry changes were minimal

We did not observe support for our hypothesis on the indirect effects of drought through litter chemistry changes. In contrast to our predictions, drought did not increase the recalcitrance of either litter type as lignin remained unchanged under drought (Figure 1b). Observations on the effect of drought on lignin have been mixed. While some studies found that litter that originated from drought environments had higher lignin than litter from ambient environments (13, 43), other studies showed that drought decreased lignin in litter of some, but not all, plant species (44). In contrast to our predictions, resource acquisition trait values generally did not change (Figures 2, 4, Table S1) whether litter chemistry changed under drought – as in grassland litter – or was unaffected by drought – as in shrub litter (Figure 1, Table S1). Previous studies have shown negative correlations between lignin fractions and decomposition rates (16–18), and lignin has also been shown to decrease decomposition rates of specific litter fractions such as cellulose and hemicellulose (33). The lack of change in lignin under drought likely contributed to a lack of change in substrate availability that explained the lack of response of resource acquisition traits to changes in litter chemistry under drought.

Drought did not have major effects on carbohydrates in the litter. While drought decreased the spectral area associated with glycosidic bonds in grass litter, grass drought litter still had more spectral area associated with glycosidic bonds than shrub litter (Figure 1d). Drought also had no effect on the carbohydrate ester band 1015-970 cm^-1^ (Figure 1h, Table S1), suggesting that drought did not decrease carbohydrate fractions in grass litter enough to influence substrate availability. Substrate availability in soil is limited by substrate diffusivity while substrate availability in litter likely is not (45), making it plausible that substrate availability in litter remains high even under low moisture conditions (46, 47). Our results suggest that grass litter chemistry might not have changed enough under drought to decrease substrate availability and investment in resource acquisition traits, while the lack of change of shrub litter chemistry under drought made it even less likely for substrate availability to change in shrub litter.

### Decomposition capabilities maintained under drought

Drought changed the composition of the grass microbial community likely through direct effects of drought stress and indirect effects of changes in litter chemistry (13, 15). The lack of a relationship between drought-induced changes in litter chemistry and resource acquisition traits despite changes in microbial community composition might indicate functional redundancy (48). Functional redundancy has been observed in soil (49) and in litter (42). Microbial community functioning tends to respond less to environmental perturbations in microbial communities with prior exposure to these perturbations (9, 50, 51). This functional resistance can be attributed to changes in community composition, such as increases in relative abundance of taxa that are less sensitive to drought (50–53) that can maintain the same function (28). While some bacterial populations that were enriched under drought in this same field experiment showed a gene-level tradeoff between drought tolerance and resource acquisition traits such that the number of CAZyme genes decreased under drought, other populations enriched under drought continued to maintain high numbers of CAZyme genes (28). Compensatory growth of functionally redundant taxa allows for microbial communities to maintain function in the face of environmental perturbations (54). Because CAZyme gene abundance for most substrates remained unchanged under drought (Figure 2, Table S1) despite changes in community composition under drought in this field experiment (13, 15), compensatory growth might have occurred as taxa that are resistant to either the direct effects of drought or drought-induced changes in litter chemistry increased in abundance to maintain decomposition capabilities.

Functional resistance to precipitation manipulations (48), as has previously been observed in a grassland (55) and a tropical rainforest (9, 52), could be another explanation for maintenance of decomposition capabilities. Repeated exposure to drought, similar to the long-term drought treatment imposed in our study, might have conditioned drought sensitive taxa to become more resistant to drought stress (52). Our results likely reflect physiological acclimation to dry conditions in such semi-arid or arid ecosystems (8, 56).

### Study limitations and future work

We specifically sampled the litter of plant species that are characteristic of the two vegetation types (grass and shrub) rather than species that are characteristic of each plot, and therefore, our litter chemistry data might only be indicative of changes in plant physiology under drought (19, 20, 43, 44) and did not account for changes in plant community composition such as shrub to grass conversions observed at our study site (21, 22). Drought has been shown to change plant community composition in observational studies over time (57) as well as in field experiments (21, 22), with drought being a factor that drives vegetation type conversion from chaparral ecosystems to exotic grasslands in California (22, 57). Such changes in plant community composition must be included in future experiments as they can change the litter that microbes decompose, affecting microbial communities, their traits, and decomposition rates (58). Furthermore, while our litter chemistry results are broadly consistent with litter chemistry results in other studies (13, 19, 20, 25–27), it is difficult to tease apart certain litter fractions and their responses to drought with our FTIR-derived litter chemistry data. A more quantitative and higher resolution method of analyzing litter chemistry changes might provide clearer results.

Consistent with our study, studies of litter decomposition in Mediterranean ecosystems so far indicate that drought-induced changes in litter chemistry either do not influence decomposition rates (13, 59) or do not influence decomposition rates as much as direct effects of drought (60). Plant litter chemistry can influence how microbial traits (8, 28) and decomposition rates (11, 12) respond to drought. Since our study indicates that microbial decomposition traits are resistant to drought-induced changes in litter chemistry following changes in plant physiology, microbial decomposition traits might be more likely to change if drought also changes plant community composition, especially if plant communities undergo type conversion.

## Materials and Methods

### Field Experiment Design

This study took place at the Loma Ridge Global Change Experiment (33°44’N, 117°42’W, 365 m elevation) near Irvine, California, USA. The climate is Mediterranean with a cool rainy season from November to April and a warm dry season from May to October of each year. The mean annual temperature is 17°C and the mean annual precipitation is 325 mm (13). We studied coastal sage scrub and grassland plots subjected to reduced or ambient precipitation treatments (22). This design led to four treatment combinations (2 vegetation types x 2 precipitation treatments). Each treatment combination had four replicate plots, for a total of 16 plots (4 treatment combinations x 4 replicate plots). The reduced precipitation treatment plots were covered with clear polyethylene tarps during a subset of winter storms, reducing annual precipitation by ∼40% (13, 22). Grassland plots (6.7 m x 9.3 m) were dominated by exotic annual grasses of the genera *Avena, Bromus, Festuca*, and *Lolium*, and forbs such as the genus *Erodium* (8). Shrub plots (18.3 m x 12.2 m) were dominated by the native shrubs *Salvia mellifera, Artemisia californica*, and *Malosma laurina* (22).

We measured decomposition rates, litter chemistry, metagenomics-derived functional gene abundance, and enzyme activity of plant litter at the field site with continued precipitation treatment (28). Plant litter was sampled on August 30, 2017, from all four replicate plots within each treatment combination. We only sampled litter from species that are representative of each vegetation type (i.e. only litter from shrub species was sampled from shrub plots of both precipitation treatments). Litter from all plots within each treatment combination was combined and mixed by hand while keeping treatment combinations separate from each other. We then made litter bags from 1 mm window-screen mesh and filled each bag with 6 g litter from one treatment combination. Litter bags were deployed on September 12, 2017, and were collected from each plot over four time points. In total, this study deployed 64 litter bags (16 plots x 4 time points), with 16 litter bags (one litter bag from each plot) being collected at each time point for laboratory analysis. We collected litter bags on November 30, 2017 (T1), April 11, 2018 (T2), November 2018 (T3), and February 2019 (T4). An aliquot of the sampled litter was ground in a coffee mixer (a quick whirl for 5 s) to create a coarse powder which was used for subsequent analyses.

### Litter Chemistry

The chemical composition of the plant litter organic matter was measured using Attenuated Total Reflection-Fourier Transform Infrared (ATR-FTIR) spectroscopy. The ground litter samples were gently pressed down on a clean surface of the germanium crystal in an ATR configuration (Smart Orbit; Thermo Fisher Scientific). Infrared light beamed from the interferometer (Nexus 870; Nicolet) was focused onto the interface between the sample and the top surface of the crystal through the lower facet. The sample spectrum was recorded with a spectral resolution of 4 cm^-1^ over the infrared range (4,000-600 cm^-1^). Data was sum-normalized before analysis. Compositional differences along the entire spectrum were studied using principal component analysis (PCA) and non-metric multidimensional scaling (NMDS) in R using the vegan package (61) with visualizations created using ggplot2 (62). Spectral ranges that showed distinct variation across the treatments as observed using PCA (Figure S1) were assigned to different functional groups for different compound classes quantified as peak area.

### Metagenomics

DNA was extracted from a 50-mg aliquot of ground litter from all 64 samples using ZymoBiomics DNA isolations kits (Zymo Research, Irvine, CA, USA) following manufacturer instructions. Sample homogenization was performed by bead beating for 5 min at the maximum speed of 6.0 m/s (FastPrep-24 High Speed Homogenizer, MP Biomedicals, Irvine, CA, USA). Gel electrophoresis, a Qubit fluorometer (LifeTechnologies, Carlsbad, CA, USA), and a Nanodrop 2000 Spectrophotometer (Thermo Scientific, USA) were used to assess the purity and concentration of extracted DNA. Library preparation and metagenomic sequencing were carried out at the University of California Davis Genome Center. We used NovaSeq (Illumina, San Diego, CA, USA) with PE150 sequencing and the default insert size of 250-400 bp. Taxonomic classification up to genus level was performed using a reads-based assessment with RefSeq database (maximum e-value cut-off of 10-5, minimum identity cut-off of 60% and minimum length of sequence alignment of 15 nucleotides) on Metagenomics Rapid Annotation using Subsystems Technology (MG-RAST) server version 4.0.3 (63).

We used Metagenome Orchestra (MAGO) (version V2.2b; 2020-03-08) (64) to produce metaSPAdes (version 3.13.0) (65) assemblies for individual samples. Within MAGO, the quality control of the paired-end reads was carried out with fastp (version v0.20.0) (66) to keep a Q30 read quality while carrying out adapter trimming. seqtk (version 1.3-r106) (67) was used to remove contigs shorter than 1,000 bp from the metaSPAdes assemblies. Contig-level data was used to assess community-aggregated functional differences across treatments. Prodigal (version 2.6.3) (68) was used to carry out gene-calling of metagenomic contigs from the individual sample assemblies which was then queried against the carbohydrate active enzymes (CAZy) database using dbCAN2 (version 2.0.11) (69). PROKKA (version 1.13.7) (70) was run in metagenome mode over the assemblies to generate respective annotations. To produce a community gene abundance table across the treatments, each dataset of quality-controlled paired-end reads was aligned against its respective assembly using BWA (version 0.7.17-r1188) (71). SAMtools (version 1.9) (72) was used to convert the alignments to binary format as well as to sort them. HTSeq (version 0.11.2) (73) was employed to count the number of reads aligned to the annotated features by PROKKA across each sample. CAZy gene abundances were normalized by total protein-coding genes predicted using Prodigal. Normalization accounts for variation in sequencing depth and assembly bias to provide absolute count data. CAZyme genes for specific substrates (cellulose, hemicellulose, polysaccharides, lignin, starch, oligosaccharides, peptidoglycan and chitin) were summed to obtain the total gene abundances linked to degradation of the substrates (74). Visualizations were made using ggplot2 (62).

### Extracellular Enzyme Assays

We performed extracellular enzyme assays on hydrolytic enzymes (Table 2) using previously reported fluorometric protocols (75, 76). Litter from each collected litter bag was homogenized in 25 mM maleate buffer with pH 6. The resulting homogenate was plated in 96-well opaque microplates with standards, controls, and serial dilutions of their respective substrates. Microplates were incubated at room temperature for four hours, and fluorescence was then measured in a plate reader. Enzyme activity was then calculated from fluorescence data (76) and divided by the dry weight of the litter that was homogenized. The resulting enzyme activity was then plotted against substrate concentration in scatterplots using *matplotlib* (version 3.3.2) in Python. The scatterplots were manually inspected for the artifact of substrate inhibition, in which enzyme activity decreases at high substrate concentrations instead of approaching V_max_ due to the substrate now acting as an inhibitor (77, 78). Leaving these data points in model fitting can underestimate V_max_ (78). These data points were removed, and the resulting enzyme activity was fitted to the Michaelis-Menten equation using the *curve_fit()* function in *scipy* (version 1.5.2) to produce V_max_ in units of µM/g dry litter/hr. V_max_ values from this curve-fitting were then subjected to further statistical analysis.

### Statistical Analysis

Additional statistical analysis, on top of PERMANOVA and PCA on litter chemistry and visualizations, was conducted in Python (version 3.8.5). Linear mixed effect models – conducted using the package *statsmodels* (version 0.12.0) – were performed on percent FTIR spectral areas of specific bands, CAZyme gene abundance, and V_max_ values with vegetation, precipitation, and their interaction as fixed effects and the collection time point – in days since deployment – and plot as random effects. Residuals were checked for normality after each model fit using the Shapiro-Wilk test from *scipy*, and the dependent variable was transformed by log_10_, reciprocal, or square root transformations and refitted until the model with the most normal residuals – having the largest Shapiro-Wilk p-value – was produced. The square root transformation was dropped as it often did not produce the model with the most normal residuals. Tukey’s pairwise comparisons were performed as a post-hoc test on levels of main effects and combinations of their interactions that were statistically significant in linear mixed effects model (p < 0.05). Cohen’s D was calculated as a measure of effect size for statistically significant main effects (Table S1).

## Author contributions

AAM, and SDA designed research; AAM coordinated the project; AAM was involved in litter sampling, experimental setup, sample processing and DNA extractions; BC performed the enzyme assays; SW and ZH performed the FTIR analysis; BC and AAM conducted all the statistical analysis and visualizations; SDA contributed reagents and analytical tools; BC drafted the manuscript with inputs from AAM and SDA; and all authors were involved in critical revision and approval of the final version.

## Acknowledgments

We thank Antonio Ribeiro for bioinformatics support; Therese Pham, Kaveh Siah, and Elizabeth Duan for supporting the enzyme activity analyses; and Jennifer Martiny and Claudia Weihe for help with experimental setup and the molecular work. We acknowledge Orange County Parks and the Irvine Ranch Conservancy for field site access.

## Data availability

The sequencing dataset generated and analyzed in the current study are available in the NCBI Sequence Read Archive through BioProject number PRJNA1178105 with accession numbers from SRR31127223 to SRR31127332 (https://www.ncbi.nlm.nih.gov/sra/PRJNA1178105).

## References

1. IPCC. 2023. Climate Change 2021 – The Physical Science Basis: Working Group I Contribution to the Sixth Assessment Report of the Intergovernmental Panel on Climate Change, 1st ed. Cambridge University Press. https://www.cambridge.org/core/product/identifier/9781009157896/type/book. Retrieved 25 August 2025.

2. Diffenbaugh NS, Swain DL, Touma D. 2015. Anthropogenic warming has increased drought risk in California. Proc Natl Acad Sci 112:3931–3936.

3. Yoon J-H, Wang S-YS, Gillies RR, Kravitz B, Hipps L, Rasch PJ. 2015. Increasing water cycle extremes in California and in relation to ENSO cycle under global warming. Nat Commun 6:8657.

4. Rapacciuolo G, Maher SP, Schneider AC, Hammond TT, Jabis MD, Walsh RE, Iknayan KJ, Walden GK, Oldfather MF, Ackerly DD, Beissinger SR. 2014. Beyond a warming fingerprint: individualistic biogeographic responses to heterogeneous climate change in California. Glob Change Biol 20:2841–2855.

5. Malik AA, Bouskill NJ. 2022. Drought impacts on microbial trait distribution and feedback to soil carbon cycling. 6. Funct Ecol 36:1442–1456.

6. Schimel JP. 2018. Life in Dry Soils: Effects of Drought on Soil Microbial Communities and Processes. Annu Rev Ecol Evol Syst 49:409–432.

7. Suseela V, Tharayil N. 2018. Decoupling the direct and indirect effects of climate on plant litter decomposition: Accounting for stress-induced modifications in plant chemistry. Glob Change Biol 24:1428–1451.

8. Malik AA, Swenson T, Weihe C, Morrison EW, Martiny JBH, Brodie EL, Northen TR, Allison SD. 2020. Drought and plant litter chemistry alter microbial gene expression and metabolite production. 9. ISME J 14:2236–2247.

9. Bouskill NJ, Wood TE, Baran R, Ye Z, Bowen BP, Lim H, Zhou J, Nostrand JDV, Nico P, Northen TR, Silver WL, Brodie EL. 2016. Belowground Response to Drought in a Tropical Forest Soil. I. Changes in Microbial Functional Potential and Metabolism. Front Microbiol 7.

10. Evans S, D. Allison S, V. Hawkes C. 2022. Microbes, memory and moisture: Predicting microbial moisture responses and their impact on carbon cycling. 6. Funct Ecol 36:1430– 1441.

11. Seres A, Kröel-Dulay G, Szakálas J, Nagy PI, Boros G, ónodi G, Kertész M, Szitár K, Mojzes A. 2022. The response of litter decomposition to extreme drought modified by plant species, plant part, and soil depth in a temperate grassland. Ecol Evol 12:e9652.

12. Santonja M, Fernandez C, Gauquelin T, Baldy V. 2015. Climate change effects on litter decomposition: intensive drought leads to a strong decrease of litter mixture interactions. Plant Soil 393:69–82.

13. Allison SD, Lu Y, Weihe C, Goulden ML, Martiny AC, Treseder KK, Martiny JBH. 2013. Microbial abundance and composition influence litter decomposition response to environmental change. 3. Ecology 94:714–725.

14. Pereira S, Burešová A, Kopecky J, Mádrová P, Aupic-Samain A, Fernandez C, Baldy V, Sagova-Mareckova M. 2019. Litter traits and rainfall reduction alter microbial litter decomposers: the evidence from three Mediterranean forests. FEMS Microbiol Ecol 95:fiz168.

15. Alster CJ, German DP, Lu Y, Allison SD. 2013. Microbial enzymatic responses to drought and to nitrogen addition in a southern California grassland. Soil Biol Biochem 64:68–79.

16. Cornwell WK, Cornelissen JHC, Amatangelo K, Dorrepaal E, Eviner VT, Godoy O, Hobbie SE, Hoorens B, Kurokawa H, Pérez-Harguindeguy N, Quested HM, Santiago LS, Wardle DA, Wright IJ, Aerts R, Allison SD, Van Bodegom P, Brovkin V, Chatain A, Callaghan TV, Díaz S, Garnier E, Gurvich DE, Kazakou E, Klein JA, Read J, Reich PB, Soudzilovskaia NA, Vaieretti MV, Westoby M. 2008. Plant species traits are the predominant control on litter decomposition rates within biomes worldwide. 10. Ecol Lett 11:1065–1071.

17. Zhang D, Hui D, Luo Y, Zhou G. 2008. Rates of litter decomposition in terrestrial ecosystems: global patterns and controlling factors. J Plant Ecol 1:85–93.

18. Bontti EE, Decant JP, Munson SM, Gathany MA, Przeszlowska A, Haddix ML, Owens S, Burke IC, Parton WJ, Harmon ME. 2009. Litter decomposition in grasslands of Central North America (US Great Plains). 5. Glob Change Biol 15:1356–1363.

19. Sardans J, Peñuelas J, Ogaya R. 2008. Drought-Induced Changes in C and N Stoichiometry in a Quercus ilex Mediterranean Forest. 5. For Sci 54:513–522.

20. Suseela V, Tharayil N, Xing B, Dukes JS. 2015. Warming and drought differentially influence the production and resorption of elemental and metabolic nitrogen pools in Quercus rubra. Glob Change Biol 21:4177–4195.

21. Ogaya R, Peñuelas J. 2021. Climate Change Effects in a Mediterranean Forest Following 21 Consecutive Years of Experimental Drought. Forests 12:306.

22. Kimball S, Goulden ML, Suding KN, Parker S. 2014. Altered water and nitrogen input shifts succession in a southern California coastal sage community. 6. Ecol Appl 24:1390–1404.

23. Madari BE, Reeves JBI, Machado PLOA, Guimarães CM, Torres E, McCarty GW. 2006. Mid- and near-infrared spectroscopic assessment of soil compositional parameters and structural indices in two Ferralsols. 1–2. Geoderma 136:245–259.

24. Zhuang J, Li M, Pu Y, Ragauskas AJ, Yoo CG. 2020. Observation of Potential Contaminants in Processed Biomass Using Fourier Transform Infrared Spectroscopy. 12. Appl Sci 10:4345.

25. Baker NR, Allison SD. 2015. Ultraviolet photodegradation facilitates microbial litter decomposition in a Mediterranean climate. Ecology 96:1994–2003.

26. Esch EH, King JY, Cleland EE. 2019. Foliar litter chemistry mediates susceptibility to UV degradation in two dominant species from a semi-arid ecosystem. Plant Soil 440:265–276.

27. Berg B, McClaugherty C. 2014. Plant Litter: Decomposition, Humus Formation, Carbon Sequestration. Springer Berlin Heidelberg, Berlin, Heidelberg. https://link.springer.com/10.1007/978-3-642-38821-7. Retrieved 25 November 2024.

28. Malik AA, Martiny JBH, Ribeiro A, Sheridan PO, Weihe C, Brodie EL, Allison SD. 2024. Bacterial population-level trade-offs between drought tolerance and resource acquisition traits impact decomposition. ISME J 18:wrae224.

29. Allison SD, Weintraub MN, Gartner TB, Waldrop MP. 2011. Evolutionary-Economic Principles as Regulators of Soil Enzyme Production and Ecosystem Function, p. 229–243. In Shukla, G, Varma, A (eds.), Soil Enzymology. Springer, Berlin, Heidelberg.

30. Allison SD, Chacon SS, German DP. 2014. Substrate concentration constraints on microbial decomposition. Soil Biol Biochem 79:43–49.

31. Chen X, Hu Y, Feng S, Rui Y, Zhang Z, He H, He X, Ge T, Wu J, Su Y. 2018. Lignin and cellulose dynamics with straw incorporation in two contrasting cropping soils. Sci Rep 8:1633.

32. Yu Z, Zhang W, He H, Li Y, Xie Z, Sailike Ah, Hao H, Tian X, Sun L, Liang Y, Fu R, Yang P. 2024. The CAZyme family regulates the changes in soil organic carbon composition during vegetation restoration in the Mu Us desert. Geoderma 452:117109.

33. Talbot JM, Treseder KK. 2012. Interactions among lignin, cellulose, and nitrogen drive litter chemistry–decay relationships. Ecology 93:345–354.

34. Bhatnagar JM, Peay KG, Treseder KK. 2018. Litter chemistry influences decomposition through activity of specific microbial functional guilds. Ecol Monogr 88:429–444.

35. Li X, Zheng Y. 2017. Lignin-enzyme interaction: Mechanism, mitigation approach, modeling, and research prospects. Biotechnol Adv 35:466–489.

36. Nakagame S, Chandra RP, Saddler JN. 2010. The effect of isolated lignins, obtained from a range of pretreated lignocellulosic substrates, on enzymatic hydrolysis. Biotechnol Bioeng 105:871–879.

37. Reith J, Mayer C. 2011. Peptidoglycan turnover and recycling in Gram-positive bacteria. Appl Microbiol Biotechnol 92:1–11.

38. Tláskal V, Voříšková J, Baldrian P. 2016. Bacterial succession on decomposing leaf litter exhibits a specific occurrence pattern of cellulolytic taxa and potential decomposers of fungal mycelia. FEMS Microbiol Ecol 92:fiw177.

39. Šnajdr J, Cajthaml T, Valášková V, Merhautová V, Petránková M, Spetz P, Leppänen K, Baldrian P. 2011. Transformation of Quercus petraea litter: successive changes in litter chemistry are reflected in differential enzyme activity and changes in the microbial community composition. FEMS Microbiol Ecol 75:291–303.

40. Romaní AM, Fischer H, Mille-Lindblom C, Tranvik LJ. 2006. Interactions of Bacteria and Fungi on Decomposing Litter: Differential Extracellular Enzyme Activities. 10. Ecology 87:2559–2569.

41. Schneider T, Keiblinger KM, Schmid E, Sterflinger-Gleixner K, Ellersdorfer G, Roschitzki B, Richter A, Eberl L, Zechmeister-Boltenstern S, Riedel K. 2012. Who is who in litter decomposition? Metaproteomics reveals major microbial players and their biogeochemical functions. 9. ISME J 6:1749–1762.

42. Matulich KL, Weihe C, Allison SD, Amend AS, Berlemont R, Goulden ML, Kimball S, Martiny AC, Martiny JB. 2015. Temporal variation overshadows the response of leaf litter microbial communities to simulated global change. 11. ISME J 9:2477–2489.

43. Zhang Z, Bodenheimer J, Scott G, Dukes JS, Suseela V. 2025. Climatic stress-induced changes in plant chemistry alter the compound-specific degradation of litter during decomposition. New Phytol 248:92–106.

44. Wilson AM, Burtis JC, Goebel M, Yavitt JB. 2022. Litter quality and decomposition responses to drought in a northeastern US deciduous forest. Oecologia 200:247–257.

45. Steinweg J, Dukes J, Paul E, Wallenstein M. 2013. Microbial responses to multi-factor climate change: effects on soil enzymes. Front Microbiol 4.

46. Manzoni S, Schimel JP, Porporato A. 2012. Responses of soil microbial communities to water stress: results from a meta-analysis. Ecology 93:930–938.

47. Schimel JP, Schaeffer SM. 2012. Microbial control over carbon cycling in soil. Front Microbiol 3.

48. Allison SD, Martiny JBH. 2008. Resistance, resilience, and redundancy in microbial communities. supplement_1. Proc Natl Acad Sci 105:11512–11519.

49. Gao W, Reed SC, Munson SM, Rui Y, Fan W, Zheng Z, Li L, Che R, Xue K, Du J, Cui X, Wang Y, Hao Y. 2021. Responses of soil extracellular enzyme activities and bacterial community composition to seasonal stages of drought in a semiarid grassland. Geoderma 401:115327.

50. Evans S, Wallenstein M. 2012. Soil microbial community response to drying and rewetting stress: does historical precipitation regime matter? 1. Biogeochemistry 109:101–116.

51. Meisner A, Snoek BL, Nesme J, Dent E, Jacquiod S, Classen AT, Priemé A. 2021. Soil microbial legacies differ following drying-rewetting and freezing-thawing cycles. ISME J 15:1207–1221.

52. Bouskill NJ, Lim HC, Borglin S, Salve R, Wood TE, Silver WL, Brodie EL. 2013. Pre-exposure to drought increases the resistance of tropical forest soil bacterial communities to extended drought. ISME J 7:384–394.

53. Evans S, Wallenstein M. 2014. Climate change alters ecological strategies of soil bacteria. Ecol Lett 17:155–164.

54. Jurburg SD, Salles JF. 2015. Functional Redundancy and Ecosystem Function — The Soil Microbiota as a Case Study, p. 29–49. In Lo, Y-H, Blanco, JA, Roy, S (eds.), Biodiversity in Ecosystems - Linking Structure and Function. InTech.

55. Hawkes CV, Shinada M, Kivlin SN. 2020. Historical climate legacies on soil respiration persist despite extreme changes in rainfall. Soil Biol Biochem 143:107752.

56. Dacal M, García-Palacios P, Asensio S, Wang J, Singh BK, Maestre FT. 2022. Climate change legacies contrastingly affect the resistance and resilience of soil microbial communities and multifunctionality to extreme drought. Funct Ecol 36:908–920.

57. Syphard AD, Brennan TJ, Rustigian-Romsos H, Keeley JE. 2022. Fire-driven vegetation type conversion in Southern California. Ecol Appl 32:e2626.

58. Liao C, Peng R, Luo Y, Zhou X, Wu X, Fang C, Chen J, Li B. 2008. Altered ecosystem carbon and nitrogen cycles by plant invasion: a meta-analysis. New Phytol 177:706–714.

59. Martiny JB, Martiny AC, Weihe C, Lu Y, Berlemont R, Brodie EL, Goulden ML, Treseder KK, Allison SD. 2017. Microbial legacies alter decomposition in response to simulated global change. 2. ISME J 11:490–499.

60. Prieto I, Almagro M, Bastida F, Querejeta JI. 2019. Altered leaf litter quality exacerbates the negative impact of climate change on decomposition. J Ecol 107:2364–2382.

61. Oksanen J, Simpson GL, Blanchet FG, Kindt R, Legendre P, Minchin PR, O’Hara RB, Solymos P, Stevens MHH, Szoecs E, Wagner H, Barbour M, Bedward M, Bolker B, Borcard D, Borman T, Carvalho G, Chirico M, Caceres MD, Durand S, Evangelista HBA, FitzJohn R, Friendly M, Furneaux B, Hannigan G, Hill MO, Lahti L, Martino C, McGlinn D, Ouellette M-H, Cunha ER, Smith T, Stier A, Braak CJFT, Weedon J. 2025. vegan: Community Ecology Package (2.7-1).

62. Wickham H, Chang W, Henry L, Pedersen TL, Takahashi K, Wilke C, Woo K, Yutani H, Dunnington D, Brand T van den, Posit, PBC. 2025. ggplot2: Create Elegant Data Visualisations Using the Grammar of Graphics (3.5.2).

63. Meyer F, Paarmann D, D’Souza M, Olson R, Glass E, Kubal M, Paczian T, Rodriguez A, Stevens R, Wilke A, Wilkening J, Edwards R. 2008. The metagenomics RAST server – a public resource for the automatic phylogenetic and functional analysis of metagenomes. BMC Bioinformatics 9:386.

64. Murovec B, Deutsch L, Stres B. 2020. Computational Framework for High-Quality Production and Large-Scale Evolutionary Analysis of Metagenome Assembled Genomes. Mol Biol Evol 37:593–598.

65. Nurk S, Meleshko D, Korobeynikov A, Pevzner PA. 2017. metaSPAdes: a new versatile metagenomic assembler. Genome Res 27:824–834.

66. Chen S, Zhou Y, Chen Y, Gu J. 2018. fastp: an ultra-fast all-in-one FASTQ preprocessor. Bioinformatics 34:i884–i890.

67. Li H. 2018. Seqtk: code walkthrough. https://lh3.github.io/2018/11/12/seqtk-code-walkthrough. Retrieved 30 August 2025.

68. Hyatt D, Chen G-L, LoCascio PF, Land ML, Larimer FW, Hauser LJ. 2010. Prodigal: prokaryotic gene recognition and translation initiation site identification. BMC Bioinformatics 11:119.

69. Zhang H, Yohe T, Huang L, Entwistle S, Wu P, Yang Z, Busk PK, Xu Y, Yin Y. 2018. dbCAN2: a meta server for automated carbohydrate-active enzyme annotation. Nucleic Acids Res 46:W95–W101.

70. Seemann T. 2014. Prokka: rapid prokaryotic genome annotation. Bioinformatics 30:2068– 2069.

71. Li H, Durbin R. 2009. Fast and accurate short read alignment with Burrows–Wheeler transform. Bioinformatics 25:1754–1760.

72. Li H, Handsaker B, Wysoker A, Fennell T, Ruan J, Homer N, Marth G, Abecasis G, Durbin R, 1000 Genome Project Data Processing Subgroup. 2009. The Sequence Alignment/Map format and SAMtools. Bioinformatics 25:2078–2079.

73. Anders S, Pyl PT, Huber W. 2015. HTSeq—a Python framework to work with high-throughput sequencing data. Bioinformatics 31:166–169.

74. Nuccio EE, Starr E, Karaoz U, Brodie EL, Zhou J, Tringe SG, Malmstrom RR, Woyke T, Banfield JF, Firestone MK, Pett-Ridge J. 2020. Niche differentiation is spatially and temporally regulated in the rhizosphere. ISME J 14:999–1014.

75. Baker NR, Allison SD. 2017. Extracellular enzyme kinetics and thermodynamics along a climate gradient in southern California. Soil Biol Biochem 114:82–92.

76. German DP, Weintraub MN, Grandy AS, Lauber CL, Rinkes ZL, Allison SD. 2011. Optimization of hydrolytic and oxidative enzyme methods for ecosystem studies. 7. Soil Biol Biochem 43:1387–1397.

77. Reed MC, Lieb A, Nijhout HF. 2010. The biological significance of substrate inhibition: A mechanism with diverse functions. 5. BioEssays 32:422–429.

78. Steen AD, Ziervogel K. 2012. Comment on the review by German et al. (2011) “Optimization of hydrolytic and oxidative enzyme methods for ecosystem studies” [Soil Biology & Biochemistry 43:1387–1397]. Soil Biol Biochem 48:196–197.

